# Pre-equilibrium biosensors: A new approach towards rapid and continuous molecular measurements

**DOI:** 10.1101/2022.03.26.485941

**Authors:** Nicolò Maganzini, Ian Thompson, Brandon Wilson, Hyongsok Tom Soh

## Abstract

Almost all biosensors that use ligand-receptor binding operate under equilibrium conditions. However, at low ligand concentrations, the equilibration with the receptor (*e.g.*, antibodies and aptamers) become slow and thus equilibrium-based biosensors are inherently limited in making measurements that are both rapid and sensitive. In this work, we provide a theoretical foundation for a novel method through which biosensors can quantitatively measure ligand concentration *before* reaching equilibrium. Rather than only measuring receptor binding at a single time-point, the pre-equilibrium approach leverages the receptor’s kinetic response to instantaneously quantify the changing ligand concentration. Importantly, analyzing the biosensor output in frequency domain, rather than in the time domain, we show the degree to which noise in the biosensor affects the accuracy of the pre-equilibrium approach. Through this analysis, we provide the conditions under which the signal-to-noise ratio of the biosensor can be maximized for a given target concentration range and rate of change. As a model, we apply our theoretical analysis to continuous insulin measurement and show that with a properly selected antibody, the pre-equilibrium approach could make the continuous tracking of physiological insulin fluctuations possible.

## INTRODUCTION

Receptor-ligand interactions are a key component of molecular biosensing technologies. Well-chosen molecular receptors confer high specificity, sensitivity, and compatibility with complex biofluids. For sensors designed to quantify low-abundance targets, high-affinity receptors are generally desirable, as they generate large binding signals at low target concentrations. However, as researchers push the limits of molecular sensitivity, bimolecular interactions encounter an inherent tradeoff between thermodynamics (*i.e.,* sensitivity) and kinetics (*i.e.,* time resolution). With high-affinity receptors and low target concentrations, it takes more time to achieve equilibrium between bound and unbound receptor states^1^. This is because the affinity of a receptor, as quantified by its dissociation constant (*K_D_*), is defined by the ratio of its off-rate to on-rate kinetics, 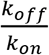 **(Figure 1a)**. The on-rate, *k_on_*, is subject to a fundamental upper limit that is determined by the physical and structural properties of the receptor and target; values range between 10^6^ – 10^7^ s^-1^M^-1^ for typical proteins, with a theoretical upper limit of ~10^8^ s^-1^M^-1^ (Refs 2–4). Consequently, differences in affinity are largely due to differences in *k_off_*, where higher receptor affinity requires a slower off-rate^2^. Most molecular biosensors are ‘endpoint’ sensors, designed to measure target concentrations in a sample of biofluid at a single point in time. In this context, slow equilibration kinetics can be remedied by allowing more time for the sensor to reach equilibrium, and thus are not a concern beyond introducing greater delay in generating a final sensor readout.

**Figure 1:**
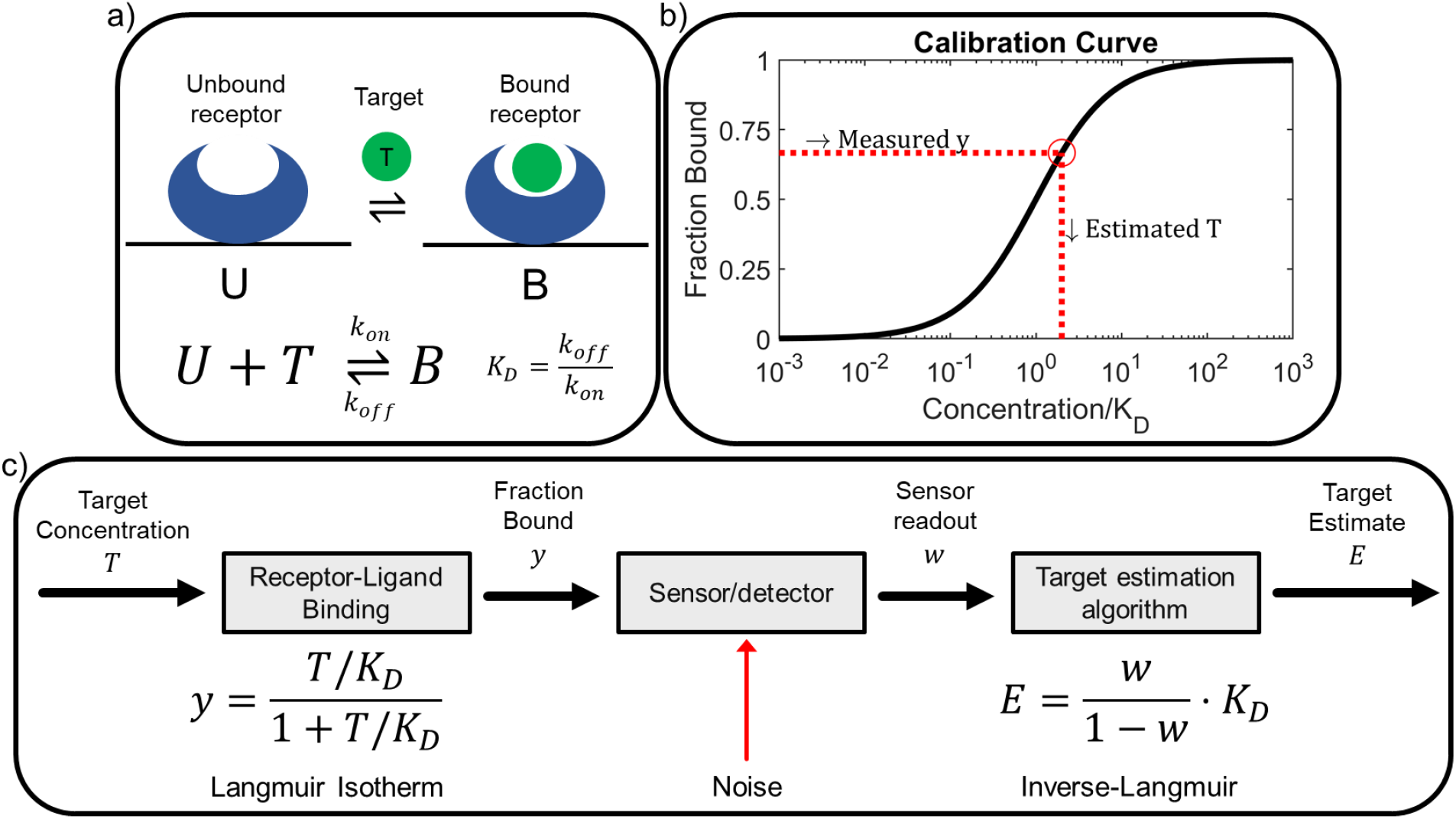
Equilibrium measurements with bimolecular sensors. **a**) A molecular receptor operating under the bimolecular assumption, where the equilibrium between unbound state U and bound state B is determined by the target concentration T. The binding properties of this receptor are described by the kinetic parameters *k_on_, k_off_*. **b**) At equilibrium, the concentration of target (*T*) is back-calculated from the measured signal representing the fraction of target-bound receptor (*y*) based on a calibration curve. **c**) A three-component model of this system, wherein *y* is related to *T* via the Langmuir isotherm. This *y* value is measured via the sensor, which also introduces noise. Finally, the target estimation algorithm (TEA) reverses the isotherm to estimate the target concentration.

More recently, “real-time biosensors” ^5–10^ that can continuously measure target concentrations have demonstrated promise as a powerful tool for biomedical applications such as drug dosing^11–20^, diagnostics^21–23^ and fundamental research^24^. In contrast with endpoint sensors, real-time sensors must adequately balance the tradeoff between thermodynamics and kinetics. Appropriate affinity is required to obtain a meaningful binding signal, but sufficiently fast equilibration kinetics are also critical to achieve reliable quantification of changing target concentrations within biologically relevant timeframes. For this reason, most real-time biosensors developed to date have targeted small-molecule analytes that are present at high concentrations, employing ~μM affinity receptors with fast kinetics, whereas real-time sensing of low-abundance analytes (such as insulin) has remained elusive. Prior efforts to address this problem have focused on increasing the rate of equilibration and tackling mass transport issues. For example, Lubken, et al.^1^ proposed a novel sensing strategy that leverages small-volume microfluidics to achieve more rapid equilibration. The same authors recently published an insightful analysis of affinity-based real-time sensors that optimizes equilibration kinetics by minimizing diffusion/advection-induced delays between bulk sample concentrations and the surface-bound receptors^25^. However, these approaches assume that equilibration is necessary for accurate quantification of target; here we question whether it is in fact necessary to reach equilibrium binding to achieve real-time analyte quantification.

In this work, we show that it is possible to achieve real-time analyte quantification with biomolecular receptors before reaching equilibrium. Instead of measuring steady state values, one can observe the equilibration dynamics of the receptor and apply a target estimation algorithm (TEA) to assess analyte concentration. In an ideal, noise-free system, this method could instantaneously determine the target concentration irrespective of the kinetics of the receptor. Noise, however, complicates the measurement of concentration changes that are faster than the kinetics of the molecular system. The sensitivity of pre-equilibrium sensors thus depends on a complex relationship between how rapidly the target concentration is changing and how rapidly the sensor can respond. Here, we investigate the accuracy of noisy biosensors that operate using our proposed pre-equilibrium sensing strategy. We first apply frequency space analysis to examine the way receptors respond to changing target concentrations at different frequencies. We find that such systems as low-pass frequency filters, wherein slowly changing concentrations (low frequencies) are tracked better than fast-changing ones (high frequencies), which is in good agreement with prior investigations^25,26^ that use similar methods. We then extend this framework to evaluate how the detector noise and the signal attenuation introduced by the molecular receptor impact the ability to reconstruct changing target concentrations. We derive design equations for pre-equilibrium sensor systems and show that the signal-to-noise ratio (SNR) in different sensing conditions can be maximized by an optimum choice of receptor kinetics. As a model, we apply our approach to demonstrate that pre-equilibrium sensing makes it possible to employ antibodies for the task of real-time insulin tracking—an application that poses a daunting challenge for equilibrium-based sensors.

## RESULTS & DISCUSSION

### Principles and limitations of equilibrium sensing

Many real-time continuous biosensors apply an equilibrium sensing technique by assuming that the concentration of target, *T*, at any point in time can be estimated from the receptor-bound fraction, *y*, at that time. For a receptor that has reached equilibrium with its surrounding sample environment, *y* will follow a concentration-dependent equilibrium binding curve determined by the receptor’s affinity, *K_D_,* which is described by the Langmuir isotherm,

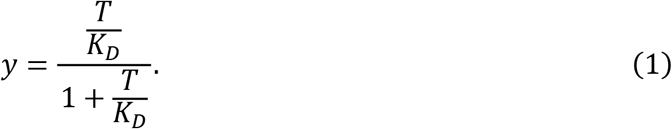

The target concentration can then be estimated from the measured signal using the inverse of this binding curve **(Figure 1b)**. Any deviations from this standard curve due to measurement noise will lead to errors in target estimation. Thus, we can describe an equilibrium sensor system as a three-component system: the molecular receptor that binds to the target, the (noisy) sensor instrument that measures the fraction of target-bound affinity reagent, and the target estimation algorithm (here, the inverse Langmuir standard curve) that back-calculates the target concentration based on this fraction (**Figure 1c**).

When this measurement technique is employed, the equilibrium assumption is critical. If measurements are performed before the receptor has reached equilibrium, the equilibrium binding curve will not accurately describe the assay results, leading to substantial errors. This places a high burden on the kinetics of the molecular receptor. The receptor must reach equilibrium at a sufficiently fast timescale compared to the changing target concentration, such that the sensor’s response directly correlates with changing molecular concentrations. The rate of equilibration, *k_eq_* = *k_on_* + *k_off_*, is bounded by fundamental on-rate limitations from diffusion and off-rate limitations imposed by the need for a very low *K_D_* to enable sensitive target detection. Thus, there are fundamental limitations to the utility of an equilibrium-based strategy for measuring rapid concentration changes in low-abundance analytes.

### Pre-equilibrium sensing

By considering the pre-equilibration dynamics of the bimolecular system, we can relax the onerous requirement that the receptor exhibits molecular kinetics that are faster than the rate of change of target concentrations. The law of mass-action describes the time-dependent behavior of a bimolecular receptor through the differential equation:

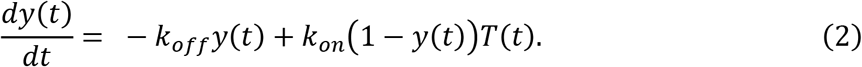

In this expression, we assume that solution-phase target molecules are in great excess relative to the sensor’s receptor, such that the solution target concentration, *T*(*t*), is unaffected by the fraction of bound receptors, *y*(*t*). By re-arranging this equation, one can determine *T*(*t*) at any time-point by measuring both the fraction of bound receptors and the rate-of-change of bound receptors, 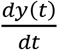, irrespective of how close or far the sensor is from equilibrium:

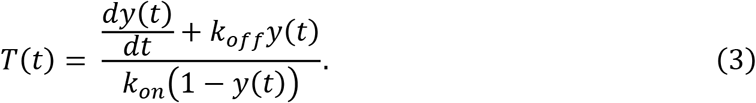

Note that when 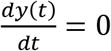 and the sensor is at equilibrium, we recover the Langmuir isotherm, Eq. 1.

Thus, in an ideal noise-free system, a receptor with any kinetic properties could be used to instantaneously determine the target concentration, irrespective of how rapidly that concentration is changing. Real-world systems are noisy, however, and this makes it more challenging to measure concentration changes that are much faster than the kinetics of the molecular system. This can be intuitively understood by considering Eq. 11 in the simplified context of two systems, a slow system (S) and a fast system (F), initially unbound with *y*(*t* = 0) = 0. The systems employee receptors with the same affinity, 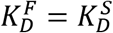, but system F’s receptor has more rapid kinetics, such that 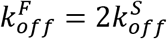 and 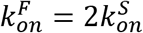 (**Figure 2a**). When exposed to a step change in concentration, the observed rate of change of the signal in F will be twice that of S. As such, the accumulated binding signal of F in a short interval of time will be twice that of S, even though both systems would tend to produce an equivalent signal at equilibrium. In a noise-free scenario, Eq. 3 would correctly reconstruct *T*(*t*) for both systems. In a real system, a fixed amount of noise would impact the slow sensor more due to its lower signal, increasing the signal-to-noise ratio and the proportion of noise in the estimated target concentration *E*(*t*). This implies that for pre-equilibrium measurement systems, sensors with slow kinetics can tolerate less noise than sensors with fast kinetics. In other words, a fast receptor will be able to measure faster-changing target concentrations more accurately than a slow receptor, given the same amount of noise. Thus, while the sensitivity of equilibrium-based sensors is only dependent on noise level and *K_D_*, the sensitivity of pre-equilibrium sensors will depend on a complex relationship between how rapidly the target concentration is changing and how rapidly the sensor can respond. This relation will in turn influence how noise propagates through the TEA, and due to the non-linearity of the Langmuir isotherm underlying receptor binding, these relationships will also depend on the thermodynamics of the system. Understanding and quantifying these relationships is essential to the successful realization and optimization of pre-equilibrium-based sensors.

**Figure 2:**
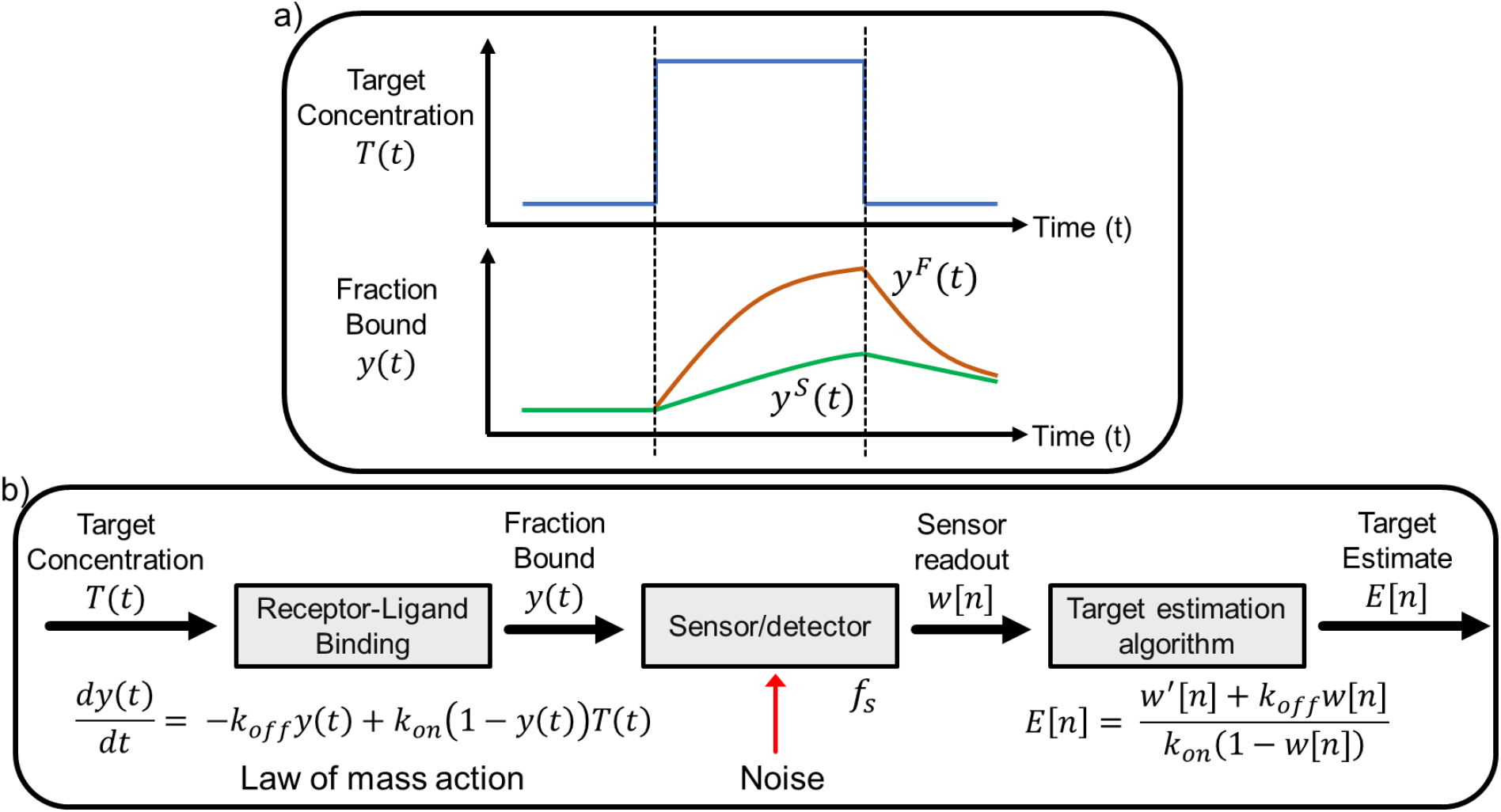
Pre-equilibrium sensing. **a**) In a short time interval, a molecular receptor with fast kinetics will accumulate more signal, *y^F^*(*t*), in response to a rapid change in target concentration, *T*(*t*), than one with slow binding kinetics, *y^S^*(*t*). **b**) An updated three-component sensor system model that incorporates pre-equilibrium equations and parameters.

Based on this understanding, we updated our initial three-component sensor system model with the pertinent pre-equilibrium sensing equations (**Figure 2b**), where the target concentration *T* and receptor-bound fraction *y* are allowed to be continuous variables in time. In this new model, the sensor samples these continuous-time signals with sampling frequency *f_s_* and generates a discrete-time sequence of measurements *w*[*n*], where the integer variable *n* replaces the continuous variable *t*. The revised TEA is the re-arranged law of mass action (Eq. 3) applied to *w*[*n*], which generates a discrete-time sequence of target estimates, *E*[*n*]. Measurement noise is shown as an additive term in *w*[*n*] introduced by the sensor, which impairs the accuracy of the TEA. In the absence of noise, this system would accurately track the target concentration regardless of receptor kinetics – we subsequently used this model to understand how measurement noise impairs accurate target estimation.

### Analysis of the pre-equilibrium sensing system is based on a frequency-domain approach

To assess our pre-equilibrium sensor model, we employed frequency-domain analysis^27^. This approach is built upon Fourier analysis, which allows us to approximate any target waveform using a sum of sinusoids of different frequencies. As we increase the number and frequency of the harmonics used in the approximation, we can capture rapidly changing features of the target waveform more accurately. **Figure 3a** shows two signals—one with slow transitions and one with sharp concentration changes—overlayed with their corresponding approximations using increasing numbers of frequency harmonics. A higher number of harmonics was required to resolve the sharper edges of the fast-changing signal. Lubken, et al.^25^, provide an analysis method that reconciles physiological concentration change rates (CCR) with corresponding frequency content for a variety of high-profile targets. By analyzing the frequency response of a receptor, we can quantify the extent to which the receptor will resolve rapid changes in the target concentration waveform. Intuitively, we expect that a receptor with fast kinetics, which rapidly reaches equilibrium, will respond to high and low frequencies in a similar manner. In contrast, the response of a receptor with slow kinetics will be impaired or attenuated at high frequencies compared to low frequencies, and thus won’t resolve the rapidly changing features of the target waveform.

**Figure 3:**
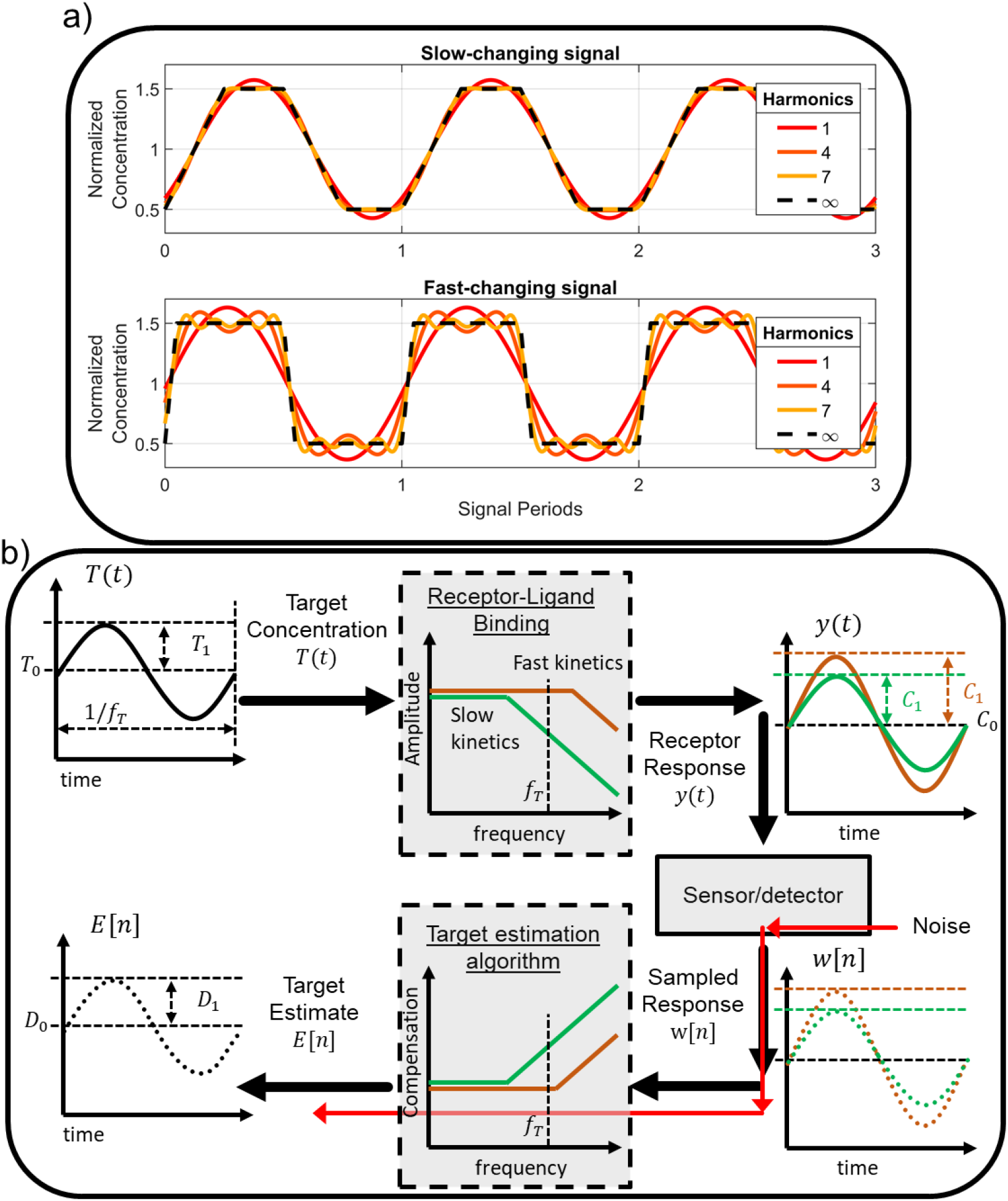
Frequency-domain analysis of a pre-equilibrium sensing system. **a**) Slow- and fast-changing target concentration signals resolved with increasingly high frequency content. High-frequency content is associated with fast-changing features in the waveform. A larger number of harmonics are required to resolve the sharp transitions of the fast-changing signal. **b**) We used a sinusoidal test signal *T*(*t*) with mean concentration *T*_0_ and amplitude *T*_1_ to analyze the receptor’s frequency-domain characteristics. A receptor with fast kinetics will respond equally well to low- and high-frequency content. A receptor with slower kinetics will attenuate high-frequency content, however, resulting in a reduced fraction of target-bound receptors, *y*(*t*). To reconstruct the original target signal *T*(*t*) from the sampled response *w*[*n*], the TEA must compensate for this attenuation. Since noise introduced by the sensor will experience this amplification as well, compensation of slow-kinetics systems will also increase the contribution of noise to the target estimate.

Through knowledge of the receptor’s kinetic parameters, one can use the TEA to estimate target concentrations by compensating for the receptor’s attenuation of high-frequency components, such that all frequency components of the target *estimate* are equal to those of the original target signal. However, this compensation will be applied to noise introduced by the sensor as well. For a receptor with slow kinetics, the algorithm must compensate for large signal attenuations, and the noise will be greatly amplified. For a fast receptor, the required amplification of high frequencies is minimal, and thus the impact of noise on the target estimate will be smaller. Thus, the limits on the amount of noise that can be tolerated in a system employing a receptor with slow kinetics will be more stringent than in a system employing a fast receptor. This means that the fast system can resolve higher frequency content than the slow system with the same amount of noise.

To carry out our frequency-domain analysis, we defined a ‘test’ target signal *T*(*t*) that comprises a sinusoidally-varying concentration of amplitude *T*_1_ summed to an average concentration, *T*_0_:

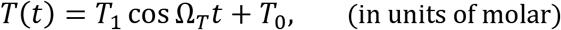

where *Ω_T_* = 2*πf_T_* is the frequency of the sinusoid in rad/s, and *f_T_* is the frequency in Hz (**Figure 3b**). We then leveraged an analysis technique termed harmonic balance to obtain mathematical relationships between *T*_0_, *T*_1_ and *C*_0_,*C*_1_ and *D*_0_, *D*_1_, where the *C* and *D* parameters respectively refer to the fraction-bound and target estimate signals, as follows:

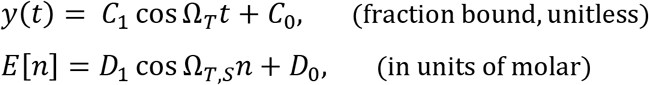

where *Ω_T,S_* = 2*πf_T_/f_S_* is the frequency of the sinusoid normalized to the sampling frequency of the detector. For example, the ratio *C*_1_/*T*_1_ quantifies the attenuation of the sinusoidal component introduced by the molecular receptor, while *D*_1_/*C*_1_ describes the compensation of the sinusoidal component carried out by the TEA.

### Frequency response of the receptor - slow kinetics act as low-pass filters

We analyze the bimolecular system in the frequency domain by applying the harmonic balance method to Eq. 1. Details of this analysis are reported in Appendix 1. The analysis results in the following relationships between *T*_0_, *T*_1_ and *C*_0_, *C*_1_.

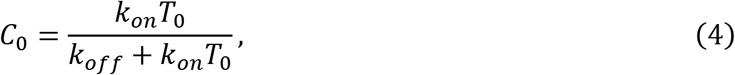

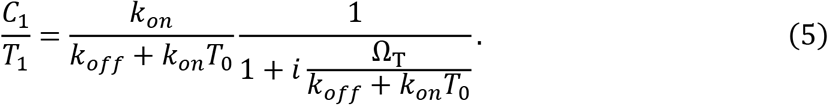

Eq. 4, which relates the zero-frequency (constant) term of *T*(*t*) to the constant term of the fraction bound signal *y*(*t*), is simply the Langmuir isotherm. This confirms that the harmonic balance analysis correctly predicts the behavior of a receptor exposed to a constant target concentration. Eq. 5 details the impact of the receptor’s kinetics on its ability to respond to target oscillations, in terms of the fraction of target-bound receptors. This equation takes the form of a first-order low-pass frequency filter with a pre-factor and describes the attenuation of the sensor response as the frequency of the target increases. The single pole present in the term has a frequency Ω_c_ = *k_off_* + *k_on_T*_0_. At this frequency, the second term becomes 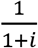, and therefore reduces the amplitude by - 3dB (~30%). This frequency is also known as the ‘corner frequency’ because at lower frequencies, *Ω_T_* < *Ω_C_*, the magnitude is largely unaffected, but for *Ω_T_* > *Ω_C_* the sensor response is subject to steep attenuation (−20 dB/10-fold attenuation per 10-fold increase in frequency). We can parametrize the corner frequency as a system-dependent ratio,

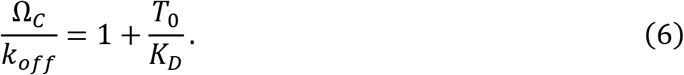

When the average concentration of target is large compared to the affinity of the sensor, the sensor operates at a high *Ω_C_/k_off_* and is capable of measuring fast-changing signals (relative to its off-rate) with minimal attenuation. This is defined as a high ‘thermodynamic operating point’ (TOP). However, when *T*_0_ ≪ *K_D_*—*i.e.,* at a low TOP—we find that *Ω_C_* ≈ *k_off_*. Thus, a fast off-rate will be required to observe rapidly changing signals with minimal attenuation.

We performed numerical simulations of a bimolecular sensor to verify agreement between the frequency-domain equations proposed above and the time-domain differential equations that describe the system (Eq. 11). We synthesized test target signals (**Figure 4a**) and numerically observed the steady state response of Eq. 11, which represents the response of an idealized receptor. We simulated three receptors with equal affinity but progressively slower kinetics, and their response is shown in **Figure 4b**. All three receptors oscillated about a mean fraction bound of *C*_0_ ≈ 0.091, which is consistent with the Langmuir relation (Eq. 1, 4) for *T*_0_/*K_D_* = 0.1. The system operating below its corner frequency (*Ω_T_/Ω_C_* = 0.1, red waveform) had the highest-amplitude response to target oscillation, while receptors with slower kinetics showed progressively smaller responses. Receptors with slower kinetics also exhibited a delayed binding response that lags changes in the target concentration, reflective of a shift in the phase of the sinusoid. Both the amplitude and phase behavior are in qualitative agreement with the analytical expression provided above (Eq. 5). We studied this further by simulating a range of systems with different kinetics and measuring their amplitude and phase response (**Figure 4c, d**).

**Figure 4:**
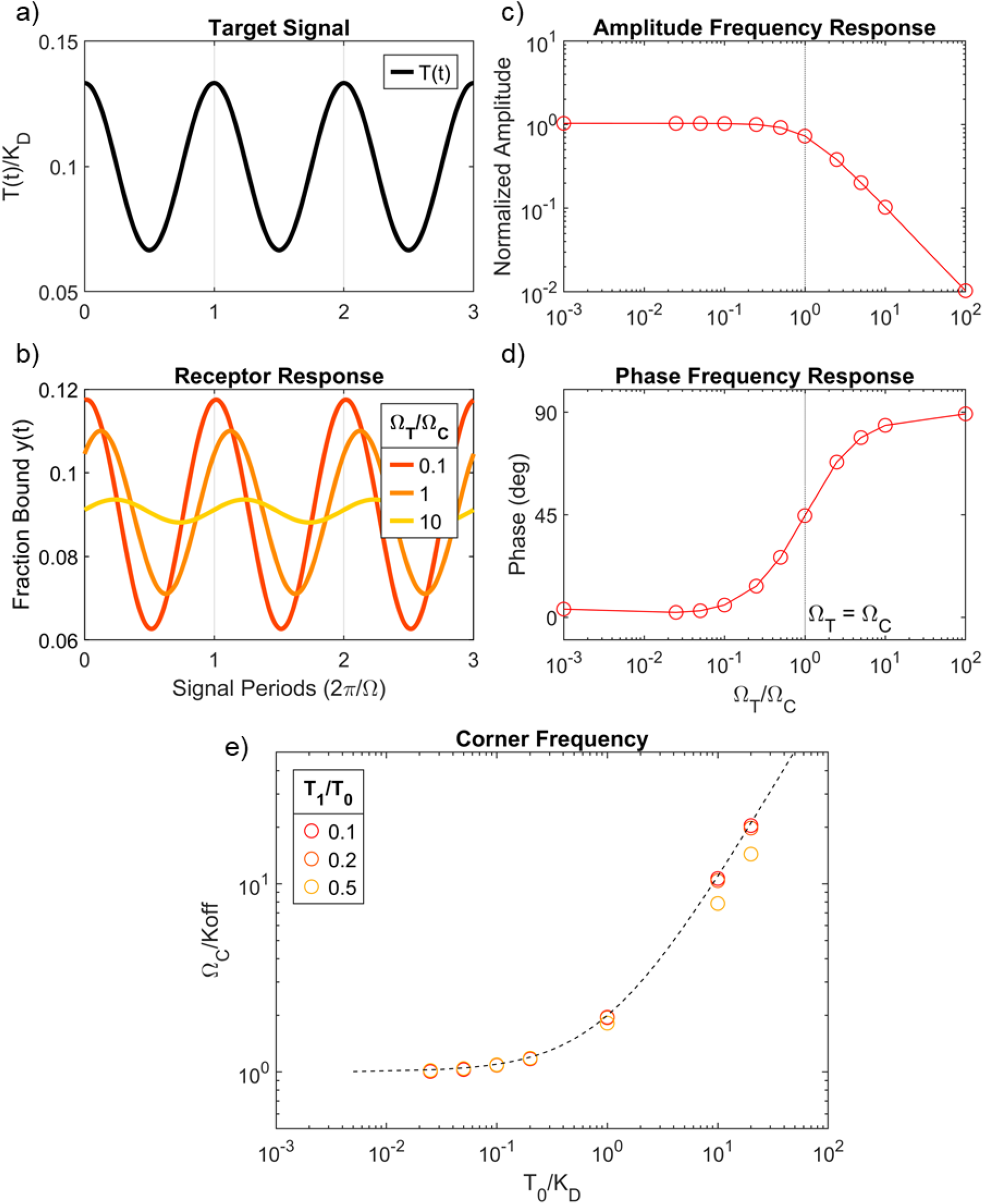
Numerical simulations of a bimolecular sensor. **a**) Example target signal employed in a simulation with *T*_0_/*K_D_* = 0.1 and *T*_1_/*T*_0_ = 1/3. Target waveforms were used as inputs to Eq. 11 to numerically simulate the response of the receptor. **b**) Simulated steady-state responses of the receptor to the target signal in **a**. Waveforms represent three different systems with progressively slower kinetics, where the ratio of target frequency (*Ω_T_*) to corner frequency (*Ω_C_*) increases from 0.1 (fastest kinetics) to 10 (slowest kinetics). **c**) Amplitude and **d**) phase measured from numerical simulations of a range of systems with progressively slower kinetics (increasing *Ω_T_/Ω_C_*). Amplitude is normalized to the expected amplitude of a system at equilibrium. Phase is measured with respect to the target waveform. The frequency response is in good agreement with a first-order low-pass filter model with corner frequency aligned at *Ω_T_/Ω_C_* = 1. **e**) Measurements of corner frequency for systems with different TOP. The predicted dependence of *Ω_C_/k_off_* on system thermodynamics (Eq. 6) was verified numerically. The dotted line represents Eq. 6, circles are measurements of *Ω_C_* (normalized to *k_off_*) from simulated systems with different *T*_0_/*K_D_*. Colors indicate target signal amplitude relative to average target concentration (*T*_1_/*T*_0_).

When target concentrations changed much more slowly than receptor kinetics (*Ω_T_/Ω_C_* ≪ 1), the measured amplitude response was very close to that expected from a bimolecular sensor operating at equilibrium. When the frequency of the target oscillations matched the corner frequency, the amplitude decreased by about 30%, and the phase increased to 45°. As target oscillation frequency increased beyond that, we saw a steep 10-fold decrease in amplitude per 10X increase in frequency. This indicates that the first-order low-pass filter model proposed above, with *Ω_C_* as its corner frequency, is a good approximation of the behavior of the receptor, as was previously reported^25,26^. Next, we verified that the proposed relation between *Ω_C_/k_off_* and the sensor TOP (*i.e.*, Eq. 6). We simulated systems with different TOPs and measured their corresponding *Ω_C_*. **Figure 4e** shows these measurements, normalized to *k_off_*, together with a plot of Eq. 6. We also simulated different amplitudes of target oscillation, *T*_1_/*T*_0_. For small oscillations, the proposed analytical expression was in good agreement with the simulations. For increasing *T*_0_/*T*_0_, however, the system deviated from the assumptions of the analytical derivation and the accuracy decreased, particularly when *T*_0_ ≫ *K_D_* and the sensor approached saturation.

The insights and equations provided in this section can be used as heuristics to evaluate the suitability of a given receptor’s performance or to estimate the kinetic requirements of a receptor, in the context of particular sensing applications. The challenge of real-time monitoring of insulin offers an interesting case study. Suppose a sensor is intended to track fluctuations in insulin levels, with an average concentration of 100 pM, and that we intend to use a receptor with a *K_D_* of 500 pM. If *k_on_* ≈ 10^6^ s^-1^M^-1^ which is typical of a “good” antibody, this receptor will have a *k_off_* ≈ 5 x 10^-4^ s^-1^. Leveraging Eq. 6, we find that *Ω_C_* ≈ 6 x 10^-4^ rad/s, which corresponds to a period of about 3 h, indicating that signals with periodic behavior faster than 3 h would experience significant attenuation by the receptor. For example, rapid changes in insulin concentration on the scale of 10 minutes, such as after a meal, would result in the sensor operating well above its corner frequency (*Ω_T_/Ω_C_* ≈ 20). This implies that changes occurring on the scale of 10 minutes would be attenuated in magnitude by approximately 20-fold compared to the response predicted by the Langmuir isotherm. Slower changes, on the scale of 1 h, would correspond to *Ω_T_/Ω_C_* ≈ 3, or an attenuation of approximately 3-fold compared to the equilibrium response. Although this poses a seemingly intractable problem in the presence of measurement noise, as described below, receptors of this type can still potentially be of value in this context.

### Frequency response of the TEA – slower kinetics result in more noise

We have observed how a receptor affects the frequency content of *T*(*t*) in the process of transducing it into *y*(*t*). Next, the sensor measures the continuous-time signal from *y*(*t*), sampling it at frequency *f_S_* to produce a discrete-time sequence *w*[*n*] with values ranging from 0 to 1. The sensor also injects noise, *N*[*n*], into *w*[*n*]. The dominant components of this noise depend on the nature of the detection modality, but we assume here that *N*[*n*] encompasses all relevant sources of random uncertainty We also make the simplifying assumption that *N*[*n*] is white gaussian noise with equal power (given by *N*_0_/2 with units of (fraction bound)^2^/Hz) at all frequencies^1^. Because electronics/detectors can operate at very rapid time-scales compared to physiological changes (*f_S_* ≫ 2*f_T_*) digital filtering or averaging techniques are typically employed to eliminate noise at frequencies higher than *f_T_*. ^28^ Since the sensor is sampling a large number of target molecules, Poisson noise is assumed to be negligible compared to detector noise. We evaluated the impact of *N*[*n*] on our ability to resolve changing molecular concentrations with slow-kinetics receptors. As shown in **Figure 3**, the frequency content of *N*[*n*] is shaped by the TEA. Since the receptor attenuates high frequencies in the signal, the TEA applies compensation by amplifying them – the slower the receptor or the higher the frequency, the greater the amplification. *N*[*n*] experiences this high-frequency amplification as well.

Thus, to accurately quantify how noise will impair target signal reconstruction, we must analyze the frequency behavior of the TEA. We applied the harmonic balance method to the equation,

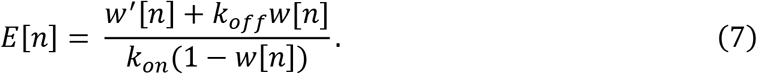

Details are reported in Appendix 2. The analysis results in the following relationships between *C*_0_, *C*_1_ and *D*_0_, *D*_1_:

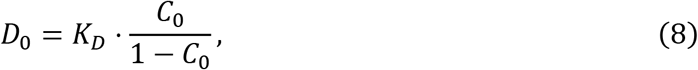

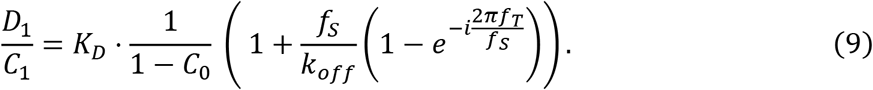

Eq. 8, which relates the zero-frequency term of *y*(*t*) to the zero-frequency term of the target estimate *E*[*n*], is simply the inverse-Langmuir. Eq. 9 describes how the TEA reconstructs the target concentration by compensating attenuated frequencies in the signal generated by the bound-receptor fraction. The equation takes the form of a discrete-time high-pass filter^28^. Considering first the limiting case of very low-frequency oscillations, *f_T_* → 0 (*i.e.,* when the sensor perfectly tracks the target oscillations), *D*_1_/*C*_1_ reduces to the frequency-independent pre-factor:

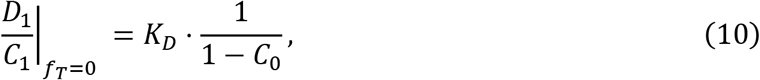

which approximates the inverse-Langmuir if *C*_1_ ≪ *C*_0_. This shows that for slow-changing signals for which the receptor reaches equilibrium, the TEA estimates target concentration simply by applying the inverse-Langmuir without further compensation. At the maximum measurable target frequency (*f_T_* = *f_S_*/2),

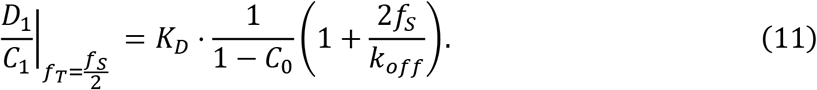

Thus, at higher frequencies, the TEA applies an extra compensation term to the inverse-Langmuir that depends on the ratio of *f_S_/k_off_*. Slow receptors with small *k_off_* will require more compensation, and therefore noise will be more amplified in such systems.

We performed simulations of the TEA to verify the agreement between the frequency-space equations proposed above and the discrete-time nonlinear differential equation that describes the TEA. We synthesized test fraction-bound signals to model the receptor response (**Figure 5a**), sampled the signals, processed them with the TEA (Eq. 7) using *K_D_* = 500 pM and *k_on_* = 10^6^ s^-1^M^-1^ as receptor parameters based on the above insulin-sensing example, and then repeated this for both a slow- and a fast-changing signal (**Figure 5b**). As expected, the amplitude of the estimated target signal was greater for the rapidly-changing signal (100–150 pM) compared to the lower-frequency signal (110–140 pM). This is because for a bimolecular sensor to generate equal fraction-bound signal changes at two different target oscillation frequencies, the change in target concentration would need to be larger in the higher-frequency system.

**Figure 5:**
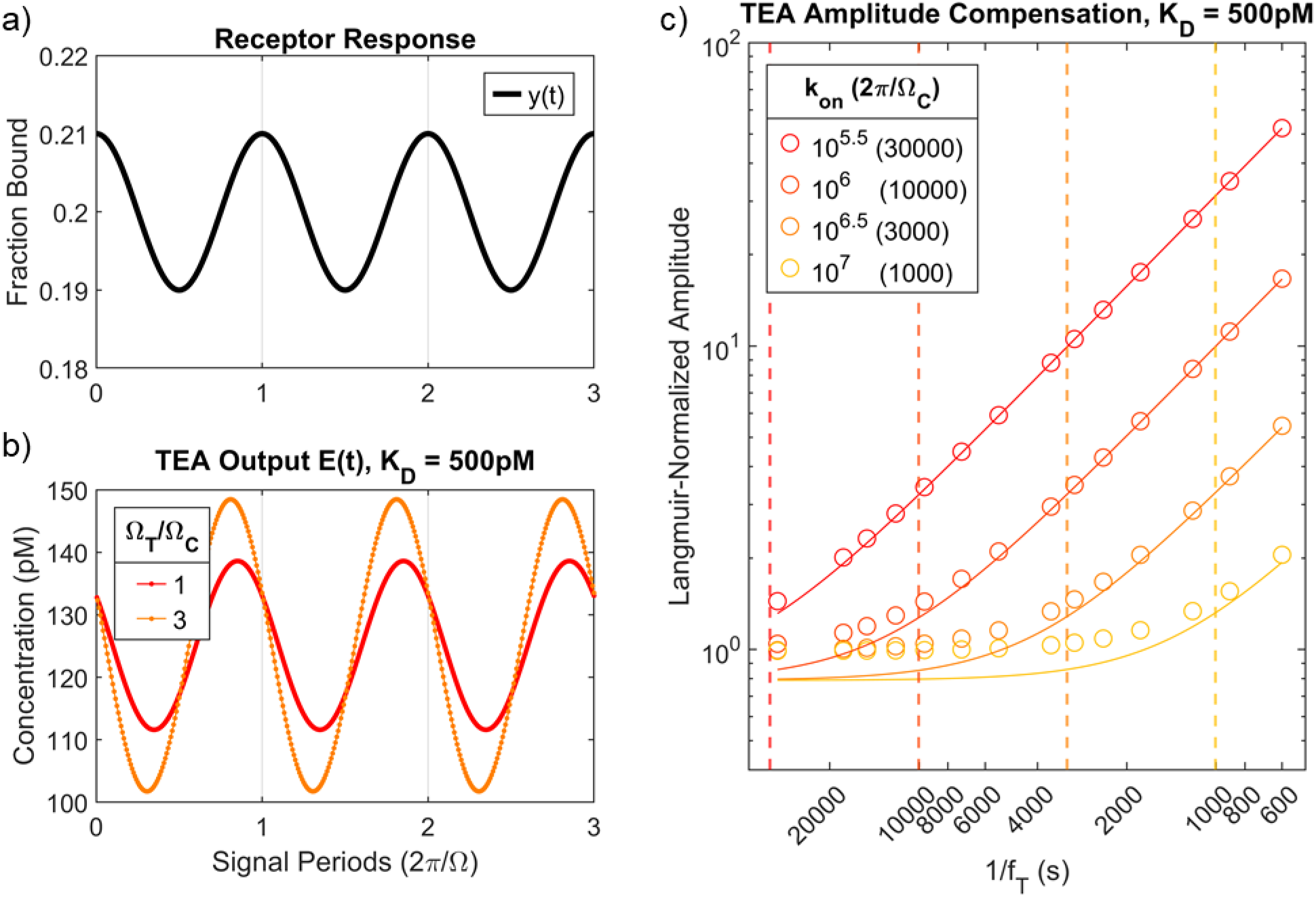
Simulations of TEA. **a**) Receptor-binding signal *y*(*t*) for a simulation with *C*_0_ = 0.2 and *C*_1_ = 0.01. **b**) The signal was sampled and processed with the TEA (Eq. 7), where the receptor’s kinetic parameters were *K_D_* = 500 pM and *k_on_* = 10^6^ s^-1^M^-1^. We performed this process with a fast-changing (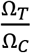; orange) and slow-changing (low *Ω_T_/Ω_C_*; red) signal to demonstrate that the TEA applies more compensation to faster-changing signals. **c**) We repeated this simulation for a range of frequencies for four different receptors with equal *K_D_* but varying *k_on_* (and thus varying *Ω_C_*). *f_S_* = 1/30 Hz. The solid lines depict Eq. 9, and circles show TEA-generated measurements of signal amplitude (Eq. 7) normalized to inverse-Langmuir predictions. Dashed vertical lines show *Ω_C_* for each receptor.

We then simulated four systems with *K_D_* = 500 pM and *k_on_* of 10^5.5^, 10^6^ 10^6.5^, or 10^7^ s^-1^M^-1^ (and thus varying *Ω_C_*) and measured the amplitude of the target estimates generated by the TEA for a range of *f_T_* values. **Figure 5c** shows this amplitude after normalization to the inverse-Langmuir equation; values >1 indicate that the TEA is compensating for attenuation introduced by the receptor. For slow-changing target concentrations, where 1/*f_T_* > 2*π/Ω_C_*, the TEA output was well-aligned with the inverse-Langmuir (i.e. normalized values close to 1). When the frequency exceeded *Ω_C_*, however, TEA compensation increases. Eq. 9 proved to be a good model for the frequency response of the TEA, as it accurately accounts for the relationship between *f_T_*, *f_S_* and *k_off_*.

We next leveraged the frequency response of the proposed TEA to obtain a closed-form expression of the amount of noise at the output of the TEA and show its dependence on receptor kinetics and *f_T_*. The average noise power at the output of the TEA, 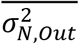, is obtained by integrating the noise power at all frequencies (details in Appendix 3). The root mean square (RMS) noise in the target estimate *E*[*n*]. which represents the average error in the target estimate, is directly related to the average noise power through 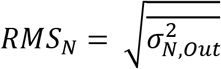, in units of molesRMS. To isolate the impact of the TEA on the amount of noise observed in *E*[*n*], we express the average noise power as:

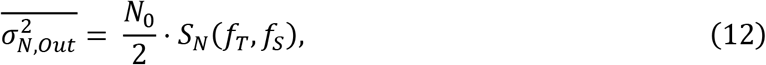

where the noise scaling function *S_N_*(*f_T_, f_S_*), with units of 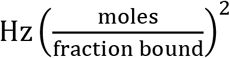, summarizes the amount of compensation applied to the white noise *N*[*n*] by the TEA. *S_N_*(*f_T_, f_S_*) is given by

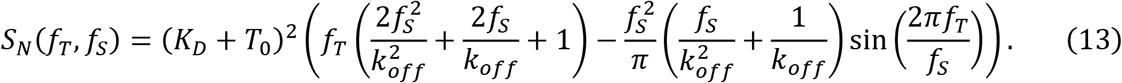

More details are provided in Appendix 3. Eq. 13 allows us to evaluate important system design decisions directly in terms of the SNR of the output target estimate *E*[*n*],

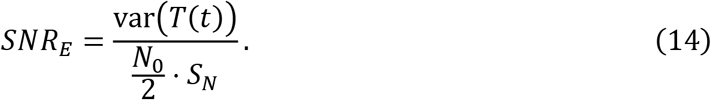

Importantly, Eq. 14 states that to improve the signal-to-noise ratio of our measurement, we must either reduce noise in our detector (lower *N*_0_/2) or reduce the noise scaling function *S_N_* by changing the sensor’s parameters.

We evaluated the analysis above by again simulating a sensor for insulin tracking. We generated a trapezoidal target signal that models the rise and fall of insulin concentrations fluctuating between 50–150 pM over a period of 10 minutes (a fundamental frequency of 1/600 Hz) (**Fig. 6a**). We simulated the response of three receptors with *K_D_* = 500 pM but progressively slower kinetics (**Fig. 6b**). We then sampled *y*(*t*) and added noise *N*[*n*] with moderate standard deviation (*σ_N_* = 0.005 fraction of bound receptor) to the samples. The resulting signal was then subjected to digital low-pass filtering. Because we aimed to resolve some of the sharper features of this signal—up to four times the fundamental frequency—we set the cutoff frequency of this filter at *f_T_* = 4/600 Hz. This filter is effectively a weighted running average of the samples, which is designed to remove unwanted frequencies. The resulting samples *w*[*n*] are shown in **Figure 6c**. Finally, *w*[*n*] was processed by the TEA to produce the target estimate *E*[*n*] (**Fig. 6d**).

**Figure 6:**
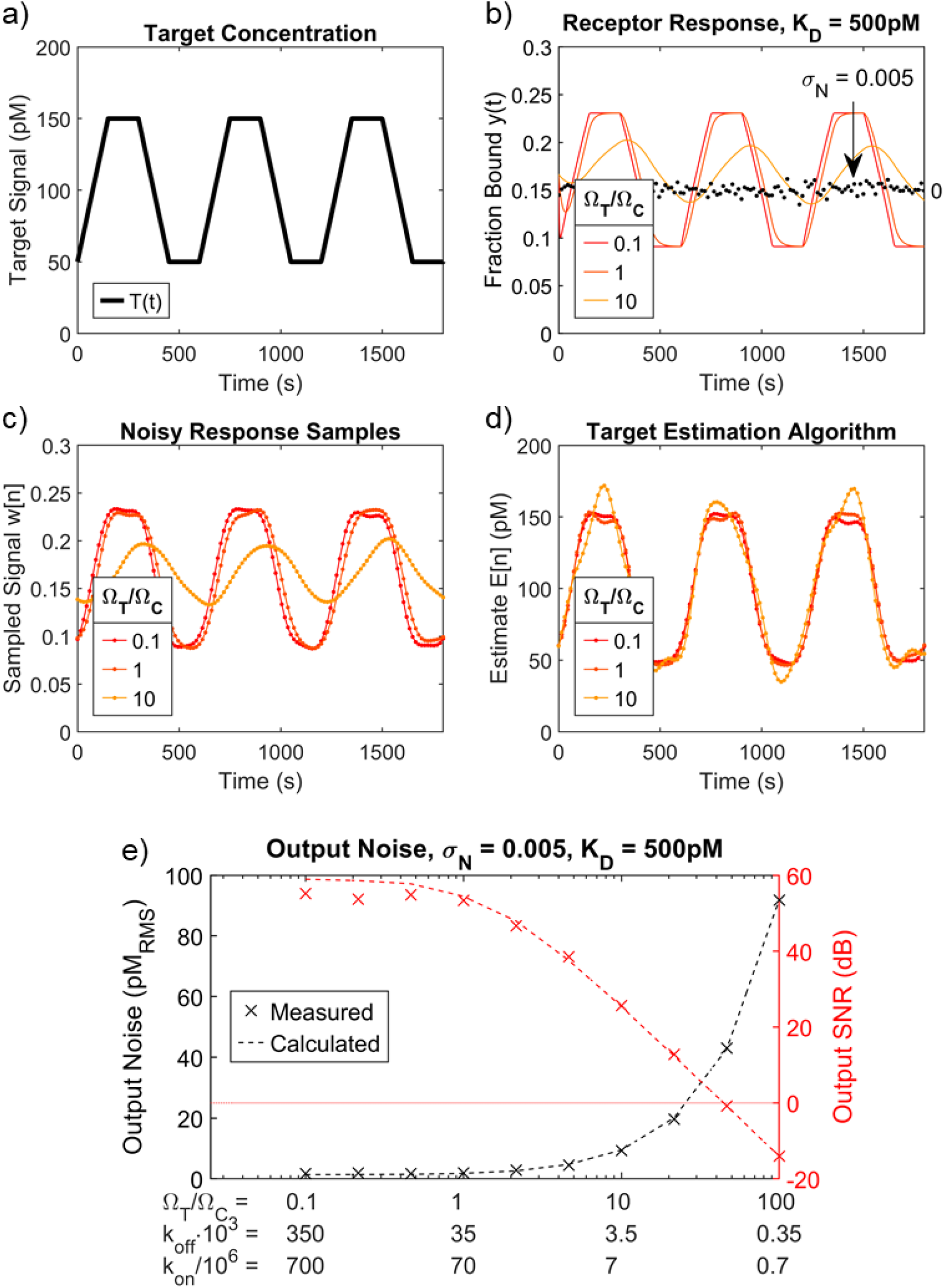
End-to-end SNR simulation of insulin sensor. **a**) A trapezoidal test concentration signal with *T*_0_ = 100 pM, *T*_1_ = 50 pM, and a period of 10 minutes (1/600 Hz). **b**) We simulated the response of three receptors with constant *K_D_* = 500 pM but decreasing kinetics, where *Ω_T_/Ω_C_* = 0.1, 1, or 10. Also shown for comparison is the noise *N*[*n*] introduced by the detector system (black dots). *N*[*n*] is zero-mean white noise with a standard deviation of 0.005 fraction bound. **c**) The receptor response was sampled with *f_S_* = 1/15 Hz, combined with *N*[*n*], and then filtered to remove all frequency content > *f_T_*. Because we aimed to resolve four harmonics above the fundamental, we choose *f_T_* = 4/600 Hz. **d**) Estimated target concentration, f⊓, at the output of the TEA. **e**) Simulations for systems with *K_D_* = 500 pM over a range of *Ω_T_/Ω_C_*, where output noise and SNR were measured. Each simulation scenario is also reported in terms of the corresponding *k_on_* and *k_off_* values. Dotted lines depict Eq. 13 and Eq. 14.

As the kinetics of the receptor became slower, the measurement was increasingly impaired by noise. The TEA applied stronger compensation to slower molecular systems, resulting in amplified noise at the output. We repeated this simulation for a wide range of receptor kinetics with a fixed *K_D_* of 500 pM and measured the output noise, defined as the error in the response of the system compared to a noiseless system (units of pM RMS), and the corresponding SNR. We compared these values with those computed through Eq. 13 and 14 (**Fig. 6e**). The results indicate that *Ω_T_/Ω_C_* < ~50 (*i.e., k_off_* > ~10^-3^ s^-1^) is required to distinguish the target signal from noise for the parameters simulated here (*i.e., T*_0_, *T*_1_, *K_D_, f_T_, σ_N_*). Importantly, the results obtained through the noise scaling function (Eq. 13) are in good agreement with the measured values, indicating that our analytical equation for *S_N_* can be used to explore the design space and optimize the system as desired.

### Optimization in terms of the noise scaling function

Considering Eq. 13 more closely, we observed that for a given *T*_0_ and *k_on_* there is an optimal choice of *k_off_* (and consequently an optimal receptor *K_D_*) that minimizes the amount of noise, and thus maximizes the SNR, of the estimated target concentration (Eq. 14). We explored this optimum for the insulin sensor case study in **Figure 7a**, where we plot the noise scaling function, *S_N_*, for two values of *k_on_*, as well as the asymptotes of *S_N_* for large and small values of *k_off_*. The location of the minimum value of *S_N_* is marked on the curves as well. As in the previous section, we used *f_T_* = 4/600 Hz and *f_S_* = 10 *f_T_*. An insulin sensor with *k_on_* = 10^6.5^ s^-1^M^-1^ would achieve optimal noise performance with a *K_D_* ≈ 1 nM (*k_off_* ≈ 3 x 10^-3^ s^-1^), while a sensor with *k_on_* = 10^7.5^ s^-1^M^-1^ would be optimized with a *K_D_* ≈ 300 pM (*k_off_* ≈ 10^-2^ s^-1^M^-1^). Deviations from these optima tend to greatly increase noise. For example, for the case of *k_on_* = 10^7.5^ s^-1^M^-1^, a receptor with a *K_D_* of 10 nM leads to 100-fold more noise than using a *K_D_* of 1 nM. The same is true for values *below* the optimal *K_D_*, showing that contrary to conventional thinking (and as demonstrated previously)^29^, a lower *K_D_* is not always better. The optimal value of *K_D_* is plotted for a range of *k_on_* and *T*_0_ in **Figure 7b**. In our analysis of *S_N_* minima corresponding to optimal *K_D_*, we noted a strong relationship between minimum *S_N_* and *k_on_* (**Fig. 7c**). For example, a 10-fold increase in *k_on_* reduced the minimum noise that could be achieved by 100-fold, indicating that the use of receptors with fast *k_on_* should offer an effective way of lessening the impact of noise on the sensor. We also note that for larger values of *T*_0_, the minimum achievable *S_N_* does not scale as directly with *k_on_*. This occurs in the regime where the optimal *K_D_* ≤ *T*_0_. The implication is that measuring small concentration fluctuations (*T*_1_) around a large mean concentration (*T*_0_) is considerably more demanding on the noise requirement than measuring the same *T*_1_ about a small *T*_0_. We also found that increasing the number of signal harmonics resolved—in this case, from 4 to 40, such that now *f_T_* = 40/600Hz (*f_S_* = 10 *f_T_*), is very costly in terms of noise performance— leads to a 1,000-fold increase in the minimum achievable noise (**Fig. 7b, c**). This is primarily because increasing the bandwidth of interest results in less filtering and more noise in the output of the TEA. It is therefore critical to choose the minimum acceptable *f_T_* that will resolve the desired features of the signal. As previously mentioned, Lubken, et al.^25^ have provided an analysis of the frequency content of several important targets based on their endogenous concentrations – their method can be used to estimate an acceptable *f_T_*.

**Figure 7:**
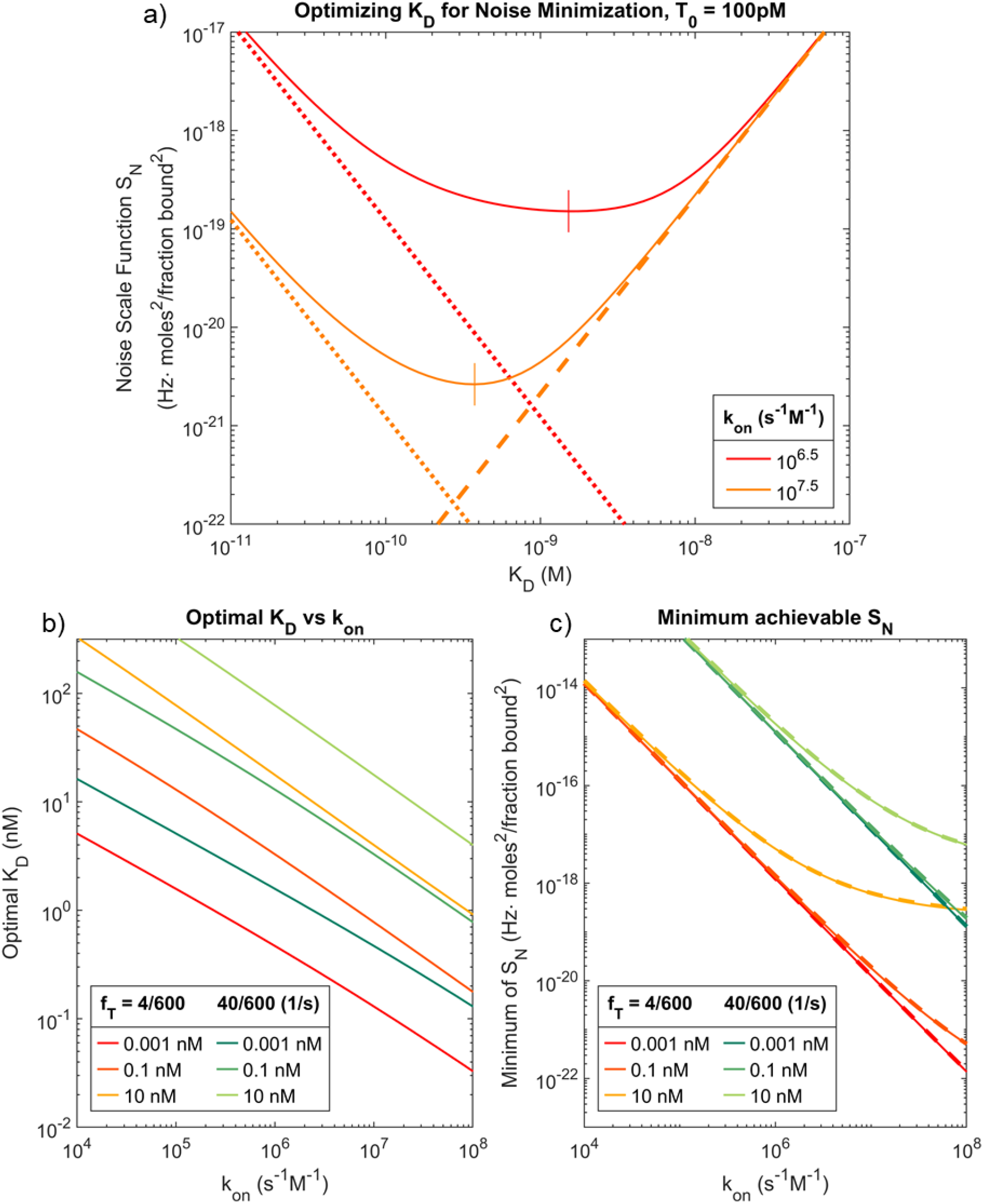
Minimizing the noise scaling function (*S_N_*). **a**) Investigating the optimal *K_D_* for two insulin receptors with *k_on_* = 10^6.5^ and *k_on_* = 10^7.5^, resolving an insulin concentration signal with *f_T_* = 4/600Hz, *f_S_* = 10*f_T_*, and *T*_0_ = 100 pM. The noise scaling function (solid lines) achieves a minimum (vertical lines) at *K_D_* ≈ 1 nM for the slow receptor and *K_D_* ≈ 300 pM for the fast one. Also plotted are the asymptotes of *S_N_* for *k_off_* → ∞ (dashed lines) and for *k_off_* → 0 (dotted lines). **b**) The optimal *K_D_* corresponding to the minimum *S_N_* was obtained for a range *k_on_* values for *T*_0_ = 0.001, 0.1, and 10 nM for insulin sensors with *f_T_* = 4/600Hz (four harmonics, target period 600 s) and *f_T_* = 40/600 Hz (40 harmonics, target period 600 s). **c**) The minimum achievable *S_N_* (corresponding to optimal *K_D_* values) for the same conditions as **b**.

As discussed above, it is critical in some cases to closely match the optimal *K_D_* in order to minimize noise. We propose a simple approximation^♥^ for the optimal *K_D_*, valid when *f_S_* ≫ *f_T_*:

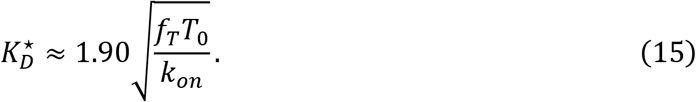

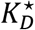 approximates the receptor affinity at which the minimum noise point is achieved and can thus be used as a rule-of-thumb for identifying optimal operating conditions. We confirmed that the *S_N_* values corresponding to 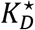 are in close agreement with the actual minimum values of *S_N_* (**Fig. 7c**), and even though 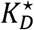 does not always accurately coincide with the optimal *K_D_*, this disparity typically occurs when the minimum in *S_N_* is shallow.

Finally, we leveraged the cumulative insights from this work to describe the design parameters of an optimal pre-equilibrium real-time insulin sensor. We maintained the target concentration parameters cited above (*T*_0_ = 100 pM, *T*_1_ = 50 pM, *f_T_* = 4/600 Hz, *f_S_* = 10*f_T_*). If we intend to design an insulin sensor with an output SNR of ~22 dB, this would correspond to an average error of about 10 pM_RMS_ for a 50 pM sinusoidal concentration signal. Based on **Figure 7**, a receptor with *k_on_* = 10^6^ s^-1^M^-1^ would perform optimally with *K_D_* ≈ 3 nM (*k_off_* ≈ 3 x 10^-3^ s^-1^) and would achieve 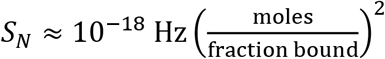. Using Eq. 12, we can calculate a noise specification for our detector: *N*_0_/2 ≤ 10^-4^ (fraction bound)^2^/Hz. This corresponds to a noise standard deviation of 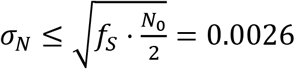 fraction bound, roughly half the one shown in **Figure 6b**. Even for the demanding sensing requirements chosen for this example, we find that the kinetic parameters are within the expected range of typical antibodies^30,31^ and that the noise requirement is tractable, based on the sensing performance of published biosensors^32–39^.This supports the idea that antibodies should be viable receptors for use in a real-time insulin sensor.

For purposes of comparison, we assessed the temporal resolution that could be achieved with a traditional real-time sensor that does not employ pre-equilibrium sensing and computes the target estimate by using the inverse-Langmuir alone. For the same insulin signal described above (*T*_0_ = 100 pM, *T*_1_ = 50 pM), an error equivalent to 10 pM_RMS_ in the target estimate is introduced when the receptor response is attenuated by ~10% of its equilibrium value. Using Eq. 5, we find that this attenuation occurs at *Ω_T_* = 1.5 x 10^-3^ rad/s or *f_T_* = 2.39 x 10^-4^ Hz. This frequency is the highest target frequency that a traditional equilibrium-based sensor could resolve with the same error as the pre-equilibrium sensor proposed above – note that this calculation is an upper-bound estimate as it neglects the impact of detector noise. Yet, the equilibrium sensor *f_T_* is ~28X slower than that of the pre-equilibrium sensor (*f_T_* = 4/600 Hz) and demonstrates that our pre-equilibrium system offers the possibility of achieving greatly improved real-time monitoring of analytes with sensors based on existing bioreceptors.

## CONCLUSION

In this work we propose a method to measure real-time changes in target concentrations prior to target-receptor equilibration. While current sensing techniques discard valuable information embedded in the binding signal prior to receptor equilibration, pre-equilibrium sensing considers the dynamics of receptor equilibration to estimate the target concentration. Frequency-space analysis of pre-equilibrium sensing shows that bimolecular target-receptor interactions can be approximated as a first-order low-pass filter, i.e. the receptor attenuates high frequency content, and smooths out rapid changes in the concentration. To account for this, we propose a pre-equilibrium target estimation algorithm that reconstructs rapid changes in target concentration by compensating for the attenuations in high frequencies introduced by slow receptor kinetics. In an ideal noise-free system, this method exactly determines the target concentration at all points in time. Noise, however, makes it more challenging to measure concentration changes that are faster than the kinetics of receptor because slower receptors require more compensation, which magnifies the noise in the target estimate. Thus, to enable informed design of pre-equilibrium systems, we derived closed-form expressions that relate parameters of the biosensor to the SNR of the estimated target concentration. We then we created numerical simulations to study the performance of the biosensor system and found good agreement between simulation and our closed-form expressions. For the case of insulin sensing, we showed how these equations can be used to determine specifications for pre-equilibrium real-time sensors. Although the equilibration rate of antibodies is typically incompatible with the timescale of physiological insulin concentration changes, pre-equilibrium sensing can be leveraged to relax this requirement.

Interestingly, we found that the SNR of the target estimate can be optimized in terms of the *K_D_* of the receptor - for a given receptor *k_on_*, there is a specific *k_off_* that minimizes the error. Perhaps counterintuitively, in many cases the optimal *K_D_* falls at significantly larger concentrations than the average target concentration *T*_0_. Conversely, a higher *k_on_* will often significantly reduce the error even when *k_off_* is left unchanged. Increasing *k_on_* is most impactful in the case of small average target concentrations, where the optimal *K_D_* falls above *T*_0_. Also, *f_T_*, i.e. the highest frequency in *T*(*t*) that the designer is interested in measuring, is an impactful design parameter that can improve performance. For example, while a physiological signal might have (small) high frequency fluctuations, these may not be of interest for the purposes of a sensing task. Choosing an *f_T_* that only encompasses features of the concentration signal that are of interest, perhaps eliminating small, high frequency features, will dramatically reduce noise requirements of the sensing system. We believe that these design principles and analytical expressions can support the implementation of pre-equilibrium-based sensing strategies. While this work has primarily studied two-state bi-molecular receptor-ligand interactions, future research could extend the theoretical insights to encompass more complex binding schemes, such as three-state receptors like aptamer switches^40^. Broadening the scope of pre-equilibrium techniques will improve biosensing performance in diverse applications by easing the requirement on receptor kinetics.

## Acknowledgements

This work was supported by the Chan-Zuckerberg Biohub, the Helmsley Trust, Bayer AG, and the National Institutes of Health (NIH, OT2OD025342). The authors would like to thank Mr. Michael Eisenstein for his editorial contributions.

## Appendix 1: Derivation of the Frequency Response of the Receptor

Consider a bimolecular sensor where U stands for unbound receptor and B is bound receptor that generates the reporting signal.

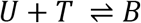

The observable sensor response is *y*(*t*) = [*B*]. The target signal is *T*(*t*), a time-varying concentration, [*T*], of target. The differential equation that describes this system is:

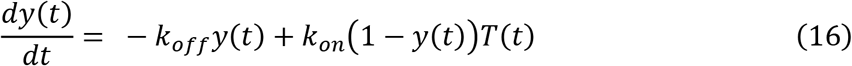

Taking the Fourier transform of the system equation, assuming no initial conditions, we obtain:

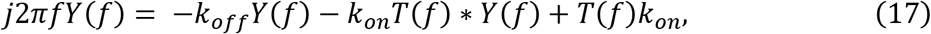

where *Y*(*f*) and *T*(*f*) represent the Fourier transforms of the sensor response and target signal, respectively, and * is the convolution operator. Specifically, we look at the case where *T*(*t*) is a sinusoidal input of one frequency *f*_T_, plus an offset term. This leads to

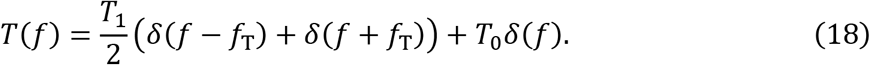

Using this expression in the convolution of Eq. 17 leads to:

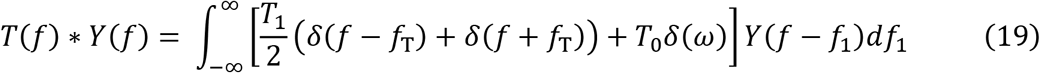

which simplifies to:

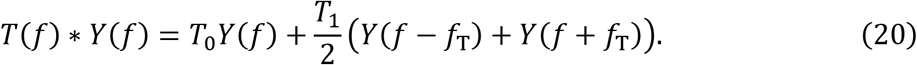

Returning to the Fourier transform of the system, Eq. 17, combining it with Eq. 18 and 20, and simplifying, we obtain:

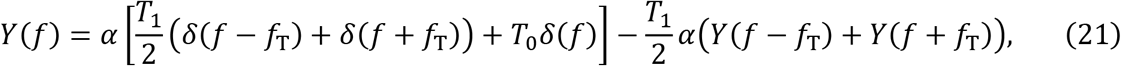

where 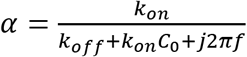. We then assume that the fraction bound signal produced by the receptor is also sinusoidal, with a frequency equal to that of the target. Thus, analogously to Eq. 18, we can write

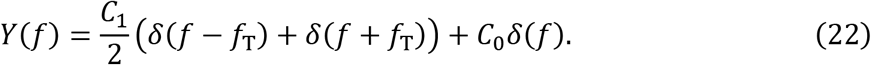

Combining Eq. 21 and *22*, we obtain an expression with multiple frequency terms, whose amplitude factors depend on *α*, *T*_0_, *T*_1_, *C*_0_, and *C*_1_. We apply harmonic balance equating all terms of the same frequency for the terms corresponding to *δ*(*f*), δ(*f* + *f_T_*) and *δ*(*f* – *f_T_*). Doing so, we obtain a system of two equations:

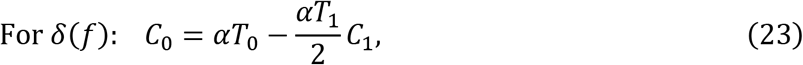

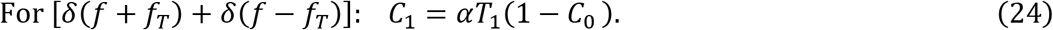

We solve these equations first by setting *f* = 0 for *C*_0_, as the harmonic corresponding to this amplitude term has zero frequency. Doing so, we obtain:

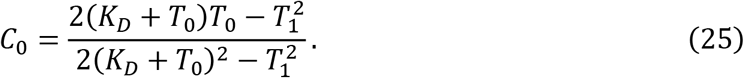

Assuming that the amplitude of oscillation is smaller than the mean concentration, *T*_1_ < *T*_0_, then Eq. 25 reduces to:

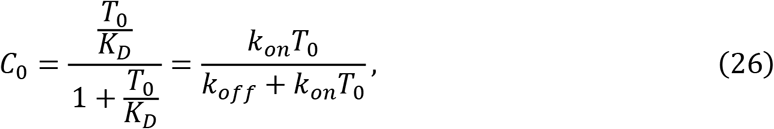

which is the Langmuir isotherm. This indicates that the harmonic balance analysis correctly predicts the behavior of the receptor responding to a zero-frequency target concentration. Next, solving the system for the transfer function *C*_1_/*T*_1_ and setting *f* = *f_T_*, we obtain:

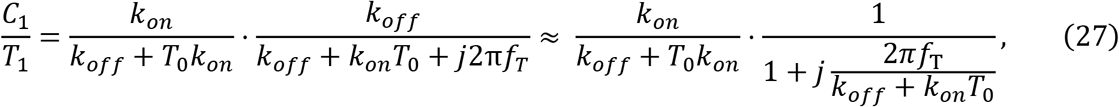

where the approximation step assumes that *K_D_* > *T*_0_, such that *k_on_T*_0_ < *k_off_*. This allows us to massage the equation into the form of a quasi-Langmuir multiplied by a first-order low-pass filter.

## Appendix 2: Derivation of Frequency Response of TEA

Re-arranging the law of mass action and solving for the target concentration signal yields:

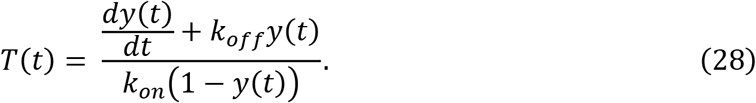

As mentioned in the main text, for the case of the model system that we analyzed, this equation is discretized to describe the operation of the TEA. This leads to the expression,

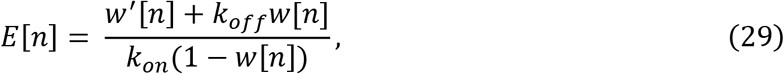

where *E*[*n*] now represents the reconstructed estimate of the target concentration. If the sampling frequency of the sensor is large compared to the frequency content of the target signal, we can estimate the derivative term *w*′[*n*] at any index *n* as:

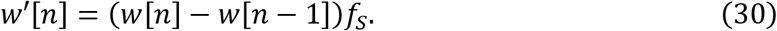

Combining Eq. 29 and 30 and taking the discrete-time Fourier transform results in the expression:

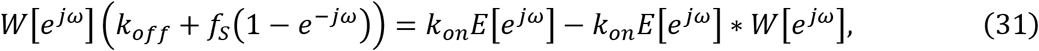

where *j* is the imaginary unit, *ω* is the normalized discrete time fourier transform frequency, *ω* = 2*πf/f_S_*, and * represents the circular convolution of the 2*π*-periodic DTFTs of *E* and *W* given by:

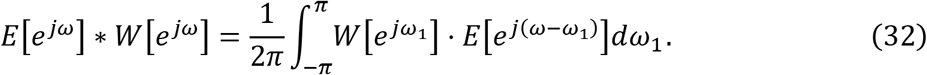

If we now assume that *w*[*n*] are samples of *C*_1_ cos(2*πf_T_t*) + *C*_0_ and *Ω_T_* = 2*πf_T_/f_S_*, then we can write the DTFT of *w*[*n*] as:

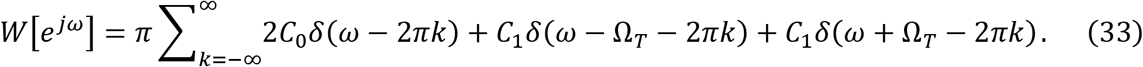

The circular convolution, Eq. 32, then becomes:

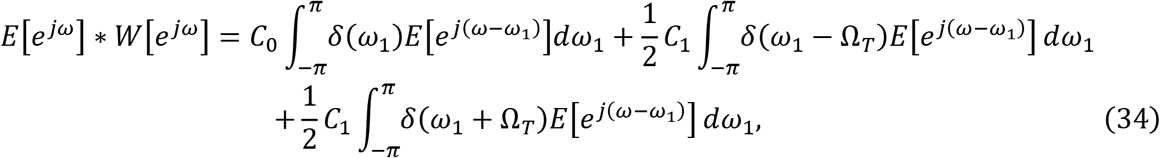

Which simplifies to:

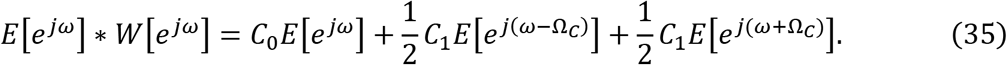

Inserting Eq. 33 and 35 into Eq. 31 and simplifying, we obtain:

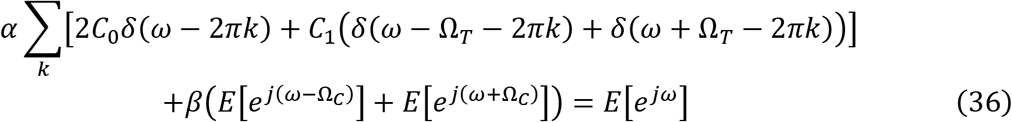

where 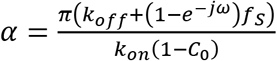 and 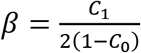. Next, we assume that the output signal *E*[*n*] will also take the form of a sinusoid of the same frequency as the input, and thus has DTFT of the form:

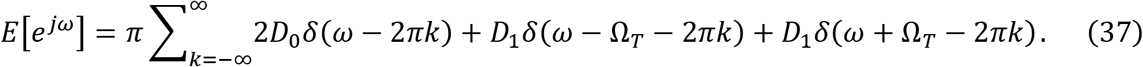

Combining Eq. 36 and 37, we obtain an equation of 2*π*-periodic DTFTs with multiple frequency components. The amplitude factors corresponding to these sinusoids depend on *α, β, C*_0_, *C*_1_, *D*_0_, and *D*_1_. We apply the harmonic balance technique, considering only the fundamental frequency terms *δ*(*ω*), *δ*(*ω* + *Ω_T_*) and *δ*(*ω* – *Ω_T_*), and equating all terms of the same frequency, we obtain the following system of equations:

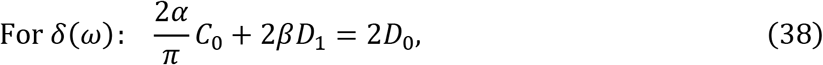

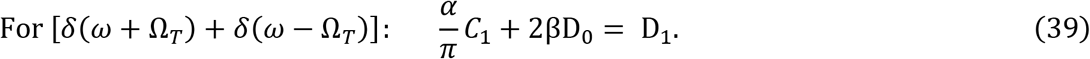

Solving these two equations for *D*_0_ first, we set *ω* = 0, as the harmonic corresponding to this term has zero frequency. Doing so, we obtain:

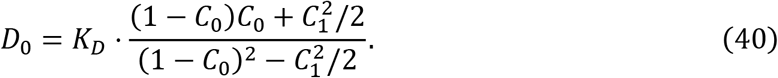

Assuming that the amplitude of oscillation is smaller than the mean concentration, *C*_1_ < *C*_0_, then Eq. 40 reduces to:

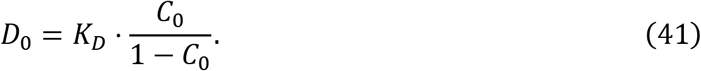

Eq. 41 is the inverse-Langmuir, indicating that the analysis correctly predicts that a constant fraction bound with value *C*_0_ at the input of the TEA will be transformed into an estimated constant concentration *D*_0_ by inverting the equilibrium behavior of the receptor.

Next, solving the system for the transfer function *D*_1_/*C*_1_, and setting *ω* = *Ω_T_*, we obtain:

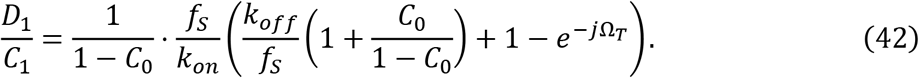

We can greatly simplify this equation by assuming that *C*_0_ ≪ 1; *i.e.,* assuming that the sensor will likely be operating in a regime where *K_D_* > *T*_0_. Taking this assumption reduces Eq. 42 to:

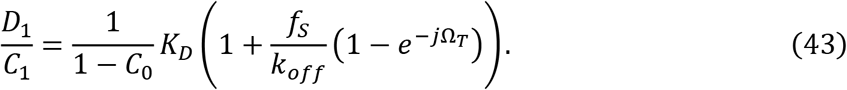

Subsequent analyses conducted in this paper can also be carried out with the more accurate expression of Eq. 42. However, if the intent is to analyze a sensor operating in the regime where *K_D_* ≪ *T*_0_, one must consider that the strong nonlinearities of the isotherm close to saturation will affect the accuracy of the analytical models presented in this discussion.

## Appendix 3: Noise Scale Function

The average noise power at the output of the TEA, 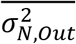, is given by the sum of output power spectral density at all frequencies. The square of the frequency response of the TEA multiplied by the power spectral density at the input of the TEA gives the output power spectral density. However, we assume that digital filtering and averaging are being used, such that all frequency content above the highest target frequency of interest, *f_T_*, is removed. Thus, the average noise power is given by:

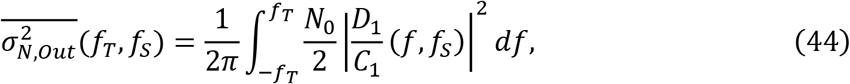

and the root mean square (RMS) noise, in units of moles_RMS_, in the target estimate *E*[*n*] is given by 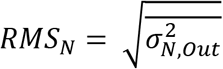. It is useful to re-write Eq. 44 as:

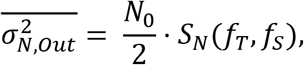

where the noise scaling function *S_N_*(*f_T_, f_S_*) with units of 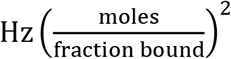 isolates the impact of the TEA. If we plug in:

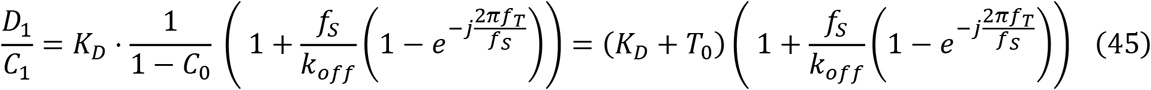

and evaluate the integral using Wolfram Mathematica, we obtain:

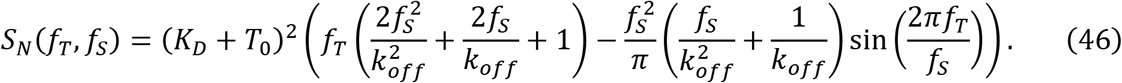

This expression for the noise scaling function can be used to calculate how the TEA propagates noise to the output. Importantly, this function can be minimized in terms of the thermodynamics and kinetics of the receptor, as is described in the main text.

## Footnotes

1 White noise is zero-mean and has power spectral density equal to *N*_0_/2 at all frequencies 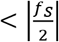.

♥ This approximation is based on the location where the asymptotes shown in Figure 7a intersect. The equations of the asymptotes are 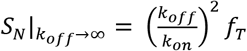 and 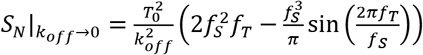. Their intersection point is 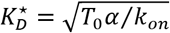, where the constant 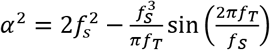. If *f_S_* ≫ *f_T_*, 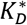 reduces to Eq. 15.

